# Cdc48/VCP and endocytosis regulate TDP-43 and FUS toxicity and turnover

**DOI:** 10.1101/668798

**Authors:** Guangbo Liu, Aaron Byrd, Fen Pei, Allison Buchanan, Eman Basha, Amanda Warner, J. Ross Buchan

**Affiliations:** Department of Molecular and Cellular Biology, University of Arizona, Tucson, AZ 85721, USA

**Keywords:** Cdc48, VCP, Endocytosis, TDP-43, FUS, ALS

## Abstract

Amyotrophic lateral sclerosis (ALS) is a fatal motor neuron degenerative disease. TDP-43 (TAR DNA-binding protein 43) and FUS (fused in sarcoma) are aggregation-prone RNA-binding proteins that in ALS can mis-localize to the cytoplasm of affected motor neuron cells, often forming cytoplasmic aggregates in the process. Such mis-localization and aggregation are implicated in ALS pathology, though the mechanisms of TDP-43 and FUS cytoplasmic toxicity remains unclear. Recently, we determined that the endocytic function aids turnover of TDP-43 and reduces TDP-43 toxicity. Here, we identified that Cdc48 and Ubx3, a Cdc48 co-factor implicated in endocytic function, regulates the turnover and toxicity of TDP-43 and FUS expressed in *S. cerevisiae*. Cdc48 physically interacts and co-localizes with TDP-43, as does VCP in ALS patient tissue. In yeast, FUS toxicity also depends strongly on endocytic function, but not autophagy under normal conditions. FUS expression also impairs endocytic function, as previously observed with TDP-43. Taken together, our data identifies a role for Cdc48/VCP and endocytosis function in regulating TDP-43 and FUS toxicity and turnover. Furthermore, endocytic dysfunction may be a common defect affecting cytoplasmic clearance of ALS aggregation-prone proteins and may represent a novel therapeutic target of promise.

## Introduction

ALS is an adult-onset neurodegenerative disease characterized by progressive motor neuron degeneration, muscle weakness and fatal paralysis due to respiratory failure, which typically occurs 3-5 years after diagnosis. ALS is mostly a sporadic disease, with ≤10% of cases exhibiting a hereditary component [1,2][3]. Many ALS-linked mutations affect proteins involved in RNA metabolism or protein clearance pathways [2], though mechanistic understanding of disease pathology remains unclear, and no effective treatment for ALS currently exists.

In ≥97% of ALS patients, a classic motor neuron hallmark of disease is re-localization from the nucleus to the cytoplasm of the RNA binding protein TDP-43, which also often forms ubiquitinated cytoplasmic protein aggregates (“inclusion bodies”) in motor-neuron cells. In approximately 1% of familial ALS patients lacking TDP-43 pathology, similar behavior is exhibited by FUS, another RNA binding protein [2]. Cytoplasmic mis-localization and aggregation of FUS is more commonly observed in another related neurodegenerative disease, Fronto-temporal lobar dementia (FTLD), where about 10% of patients exhibit FUS aggregates. The remaining 90% of FTLD patients exhibit either aggregates of TDP-43 (45%) or the microtubule-associated protein Tau (45%) [2]. TDP-43 and FUS possess similar domains including RNA recognition motifs (RRMs) glycine rich domains and/or prion-like domains, and have been proposed to affect multiple steps of mRNA metabolism [4][5].

Under conditions of cellular stress, both TDP-43 and FUS re-localize into cytoplasmic stress granules (SGs) [6,7], which are dynamic mRNA-protein assemblies implicated in regulating mRNA function[8,9]. Given this, and in light of ALS-associated mutations that generally reduce SG dynamics, SGs have been theorized to promote formation of TDP-43 and FUS aggregates that are observed in ALS patients [10,11], though recent studies suggest SG-independent aggregation mechanism also likely exist [12][13][14]. Regardless, various studies indicate that TDP-43 and FUS aggregates, or simply excessive cytoplasmic localization of TDP-43 or FUS, may result in a toxic gain of function that leads to motor-neuron degeneration [15–18][19][20,21]. While the mechanism of such toxicity remains unclear, preventing aggregation and/or enhancing clearance of cytoplasmic TDP-43 and FUS is of considerable therapeutic interest. This approach has shown promise in various ALS models where preventing SG assembly or upregulating cytoplasmic protein turnover has been tested [22–26].

Recently, we and others demonstrated that under normal growth conditions, TDP-43 toxicity, aggregation and protein turnover in yeast and human cell line models depends strongly on endocytic function [26,27]; such effects were largely independent of autophagic or proteasomal contributions. Defects in endocytic function increased TDP-43 protein levels, aggregation and toxicity, whereas genetically enhancing endocytosis rates suppressed these phenotypes. Defects in endocytic function also exacerbated motor neuron dysfunction in an ALS TDP-43 fly model, whereas endocytic enhancement suppressed such dysfunction [26]. Finally, expression of cytoplasmic TDP-43, particularly aggregation prone forms in human cells, correlated with impairment in endocytosis function itself [26]. Key remaining questions include better defining how endocytosis-dependent TDP-43 clearance is achieved, how cytoplasmic TDP-43 inhibits endocytosis, and whether endocytic inhibition is observed with other genetic ALS models.

Cdc48, and its human homolog Valosin-containing protein (VCP/p97), is a type II AAA+ ATPase chaperone that acts as a “segregase” of ubiquitinated proteins. Cdc48/VCP removes ubiquitinated proteins from complexes or membranes, and often (though not always) aids targeting of said proteins for degradation by autophagic or proteasomal means [28] [29]. Cdc48/VCP functions in many diverse cellular processes including the DNA damage response, cell cycle control, autophagy, proteolytic turnover, endocytosis and SG clearance [29–32][33]. Specificity for Cdc48/VCP function is typically derived from interactions with various cofactors that help recruit Cdc48/VCP to distinct ubiquitinated substrates [28,29].

VCP is mutated in a small fraction of ALS patients (1% of sporadic and familial ALS), and in most patients with Inclusion Body Myopathy with early onset Pagets disease and frontotemporal dementia (IBMPFD) [34,35]. As with ALS, IBMPFD also commonly exhibits cytoplasmic TDP-43 aggregates in affected cells (e.g. muscle cells, frontal cortex neurons) [34,35]. Additionally, neurodegenerative effects of VCP mutants in flies depends upon TDP-43 cytoplasmic aggregation [36]. Finally, disease-alleles of VCP lead to accumulation of cytoplasmic TDP-43 aggregates in human cells [32] and mouse models [37]. Given these observations, we were curious to define the link between Cdc48/VCP function and accumulation of pathological protein aggregates in ALS, FTLD and IBMPFD, focusing on if Cdc48/VCP aids TDP-43 and FUS turnover, and if so by what mechanism.

Several studies have shown strong links between cytoplasmic TDP-43 and FUS protein localization, aggregation and neuron loss [15,16][38]. However, the mechanism of how cells clear these apparently toxic cytoplasmic aggregates is still unclear. In this report, we demonstrate that like TDP-43, cytoplasmic FUS expression impairs endocytosis rates in yeast. Interestingly, inhibition of Cdc48 and Ubx3 (a Cdc48 endocytosis-promoting co-factor) leads to defects in FUS and TDP-43 turnover and increased toxicity, while inhibition of VCP causes accumulation of TDP-43 and FUS protein and increases TDP-43 cytoplasmic aggregation. Additionally, Cdc48 physically interacts with and co-localizes with TDP-43 and endocytic proteins. Finally, TDP-43 cytoplasmic aggregates co-localize with VCP in ALS patient tissue. Taken together, these data suggest a role for Cdc48/VCP-facilitated endocytosis in regulating TDP-43 and FUS turnover and thus toxicity, which may represent a novel therapeutic target for numerous devastating neurodegenerative diseases characterized by TDP-43 and FUS cytoplasmic pathology.

## Materials and methods

### Yeast strains and growth conditions

Untransformed yeast strains were cultured at 30°C with YPD medium. Strains transformed with plasmids were grown in SD medium with appropriate nutrients for plasmid selection. In liquid culture, strains expressing *GAL1*-driven TDP-43 and FUS from plasmids (or vector controls) were initially cultured overnight in either 2% sucrose (if starter cultures were to be used for spotting assays) or 0.25% Galactose and 1.75% Sucrose (if starters were to be used for microscopy assays), then back diluted to OD600 0.1 in identical media, followed by growth to mid-log (OD600 0.3-0.6). 0.25% Galactose, rather than 2%, ensures more modest expression and toxicity of TDP-43 [26] and FUS (not shown), which facilitates identification of enhancer mutants. Transformation was performed by a standard LiAc method. All strains and plasmids used in this study are listed in Table S1.

### Yeast serial dilution assays

Yeast transformed with TDP-43, FUS or empty vector plasmids were grown overnight as described above, back-diluted to an OD_600_ of 0.2, serially diluted as indicated in figure legends, and spotted onto identical agar media.

### HEK293A and TDP-43-GFP lentiviral integrated cell culture

HEK293A cell culture and generation of TDP-43-GFP stable integrated lines has been previously described[26]. Briefly, HEK293A cells with lentiviral integrated TDP-43-GFP or TDP-43 made to express a 35kDa TDP-43 fragment (“TDP-35-GFP”; amino acids 90-414; a cytoplasmic aggregation-prone allele mimicking cleavage by caspases [39]) were cultured at 37°C in DMEM medium (HyClone, SH30022.01) with 10% FBS (Gibco, 26140079), 50 U/ml penicillin and 50 μg/ml streptomycin (Gibco, 15140-122). DbEQ (Apexbio Technology, A862910), a VCP inhibitor, was added at a concentration of 10uM for 1 hour prior to live-cell Deltavision microscopy (see below).

### Fluorescence Microscopy

The methods have been described in detail previously [32]. Briefly, early stationary phase (OD_600_ ≥3.0) or mid-log cells (OD_600_ 0.3-0.6) cells were examined using a DeltaVision Elite microscope. Data was analyzed by Fiji [40]. At least 50 cells (yeast or human) were analyzed per biological replicate, and all shown images are representative of a minimum of 3 biological replicates. % value indicates co-localization of TDP-43 or FUS with indicated proteins, which in yeast was scored in a blind manner with a minimum of 50 cells per biological replicate.

### Western Blotting and protein stability assays

Western blotting was conducted as described previously [41]. Primary antibodies were as follows: α-GFP (902602; Biolegend), α-TDP-43 (60019-2-Ig; ProteinTech), α-Vinculin (V4505; Sigma-Aldrich), α-Pgk1 (ab113687; Abcam), α-VCP (ab11433; Abcam), α-TAP (Thermofisher CAB1001) and α-GAPDH (MA5-15738, Invitrogen). For TDP-43 and FUS protein stability assays in yeast, expression was halted either by addition of cycloheximide at 0.2mg/ml (translational shut-off) or additional of 2% glucose to the growth media (transcriptional shut-off), as indicated in figure legends.

### Endocytosis rate assays

Mid-log yeast cells (OD_600_ 0.3-0.6) were harvested and incubated on ice for 10 minutes, then suspended in YPD containing 8μM FM4-64 (T3166, Thermo Fisher Scientific) at 25°C and imaged at indicated time periods. HEK293A cells were kept on ice for 10 minutes, washed with live cell imaging solution (Thermo Fisher Scientific, A14291DJ) and then incubated with 25µg/ml fluorescent transferrin (T23362, Thermo Fisher Scientific) at 37°C for 15 minutes. Cells were then washed again with live cell imaging solution, imaged and quantified for intracellular fluorescence using Fiji.

### Patient tissue immunofluorescence

Samples were dewaxed by xylene and fixed with successive incubations in 100%, 100%, 90%, 80%, 70% ethanol and distilled water for 10 minutes. Antigen was retrieved by keeping tissue slices in boiled sodium citrate buffer for 20 minutes. Endogenous peroxidases were quenched with 3% H_2_O_2_ for 30 minutes. Each slice was then blocked with 5% goat serum for 1 hour and then incubated with primary antibody overnight at 4°C. Following 4 TBS-T washes, each slice was incubated with an appropriate alexa fluor conjugated secondary antibody (Thermo Fisher Scientific). Slices were then washed 4 times in TBS-T, mounted with ProLong Gold Antifade DAPI mounting media (Molecular Probes, P36931) to stain the nuclei, and imaged.

### Immunoprecipitation analyses

10 ml cultures of appropriately transformed yeast with suitable selective SD media were grown to mid-log growth phase (0.3-0.6 OD600), cell pellets harvested with 13000g centrifugation at 4°C, and resuspended in 1ml of RIPA cell lysis buffer. 50% volume of glass beads (BioSpec Products, 11079105) were added, and cells vortexed for 20s before being incubated on ice; this was repeated twice more. Cells were centrifuged at 13,000g for 15mins to separate unylsed cells and insoluble cell debris. Supernatant was harvested, and protein concentration assessed for normalization via Bradford assay. Supernatants were incubated with magnetic dynabeads (Thermo fisher scientific) labelled with either αTAP antibody or αGFP antibody for 2 hours at 4C, followed by three bead washes in lysis buffer. Bound proteins were eluted with gel loading buffer, boiled in a standard SDS gel loading buffer at 95C for 5 minutes, and then subject to standard Western blot analyses.

## Results

### Cdc48 co-localizes with and mediates TDP-43 degradation and toxicity

VCP disease alleles are associated with cytoplasmic re-localization and aggregation of TDP-43 [32,36,37], and mutant VCP toxicity in a fly model of ALS depends upon the cytoplasmic accumulation of TDP-43 [36]. However, mechanistic details of why VCP impairment affects cytoplasmic TDP-43 accumulation remain unclear. We began to address this using an established yeast TDP-43 model [18], focusing on interactions with the yeast VCP homolog Cdc48.

First, the localization of TDP-43 and Cdc48 was examined. Interestingly, expression of TDP-43 induced re-localization of Cdc48 from a predominantly nuclear region (mostly peri-nuclear in mid-log cells) into cytoplasmic foci that exhibited approximately 60% co-localization with TDP-43 foci in mid-log and stationary phase conditions (Fig. 1A). Consistent with co-localization in foci, reciprocal immunoprecipitation of Cdc48-TAP and TDP-43-YFP revealed a robust interaction between Cdc48 and TDP-43 (Fig. 1B-C); this is consistent with prior VCP-TDP-43 co-immunoprecipitation data from human cell culture and brain tissue [42]. In yeast, TDP-43 cytoplasmic foci co-localize with SGs, as has been previously shown [43], and which we have also confirmed with various stress granule marker proteins (data not shown). Consistent with this, Cdc48-TAP immunoprecipitation also revealed a robust interaction with Pab1-GFP, a core SG component (Fig. 1D). Taken together, our data suggests that Cdc48 co-localizes with and physically interacts with TDP-43 and Pab1, likely at least partially within a SG context.

**Fig 1.**
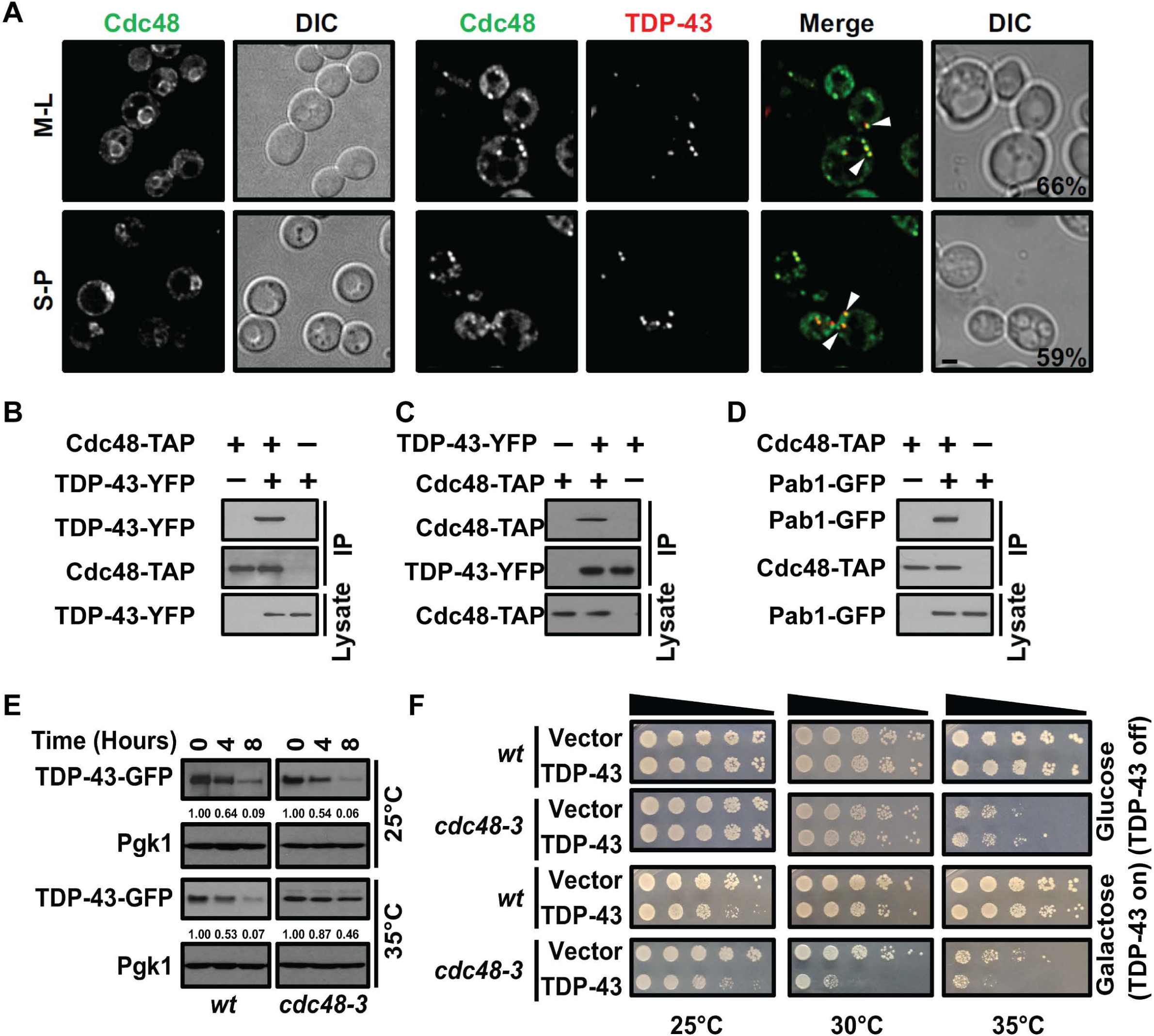
Cdc48 interacts with TDP-43 and regulates TDP-43 turnover and toxicity. (A) Cdc48-GFP strain alone (left), and Cdc48-GFP strain transformed with a TDP-43-mRuby2 plasmid (right) were imaged in both mid log (M-L) and stationary phase (S-P). Arrowhead and percentage values indicate Cdc48 and TDP-43 co-localization. Scale bar: 2 µm. (B) Cdc48-TAP strain was transformed with an empty vector or TDP-43-YFP plasmid, and then TAP immunoprecipitation was performed. (C) As in (B), but with GFP immunoprecipitation i.e. reciprocal pull down. (D) Cdc48-TAP strain with TDP-43 was transformed with vector or Pab1-GFP and the interaction was determined. (E) WT and *cdc48-3* strains were treated with 0.2 mg/ml Cycloheximide (CHX) for the indicated time at either 25°C or 35°C. TDP-43 protein levels detected and normalized relative to a PGK1 loading control. (F) Serial dilution growth assay of WT and *cdc48-3* transformed with empty vector or *GAL1*-regulated TDP-43-YFP plasmid at different temperatures.

Next, we tested whether Cdc48 affected TDP-43 stability given the many roles Cdc48 plays in protein turnover. Since Cdc48 is an essential gene, we turned to a commonly used temperature sensitive (*ts*) mutant, *cdc48-3*, which possesses two missense point mutations, P257L and R387K that inhibit Cdc48’s ATPase activity [44]. Protein degradation rate was measured by adding Cycloheximide (CHX) to block new protein synthesis. At the permissive temperature of 25°C, no change in TDP-43 turnover rate was examined in WT or *cdc48-3* strains. However, at 35°C, which strongly impairs Cdc48-3 function, TDP-43 turnover rates were significantly decreased in the mutant strain relative to WT (Fig. 1E). Finally, TDP-43 toxicity in yeast, evidenced by reduced cell growth in TDP-43 transformants relative to empty-vector control strains [10], was exacerbated by inactivation of Cdc48 (Fig. 1F). Specifically, this was evident at temperatures that modestly (30°C) and strongly (35°C) inhibit Cdc48-3 function. Together, these data demonstrate that Cdc48 regulates TDP-43 protein stability and toxicity in yeast.

### Cdc48 regulates TDP-43 toxicity in part by endocytosis

Cdc48 specificity of function is largely determined by use of co-factor proteins that help recruit it to its targets. In yeast, the Ubx family of proteins are a well described example of such proteins [45]. Thus, we first examined TDP-43 toxicity in a range of Ubx null strains (Fig. 2A). Importantly, only deletion of *UBX3*, which functions along with Cdc48 in yeast endocytosis [46], clearly exhibited enhanced TDP-43 toxicity (Fig. 2A). No clear effect with any other Ubx mutants on TDP-43 toxicity was observed, including *UBX1* (*SHP1*) or *UBX2* deletion, which facilitate Cdc48 function in autophagy and proteasomal turnover [47,48], (Fig. 2A). These findings are consistent with our prior work that endocytosis plays a greater role than autophagy in regulating TDP-43 toxicity and turnover in yeast during normal growth [26]. Finally, like Cdc48, Ubx3 foci localized with TDP-43 foci at a frequency of 53% (Fig. 2B).

**Fig 2.**
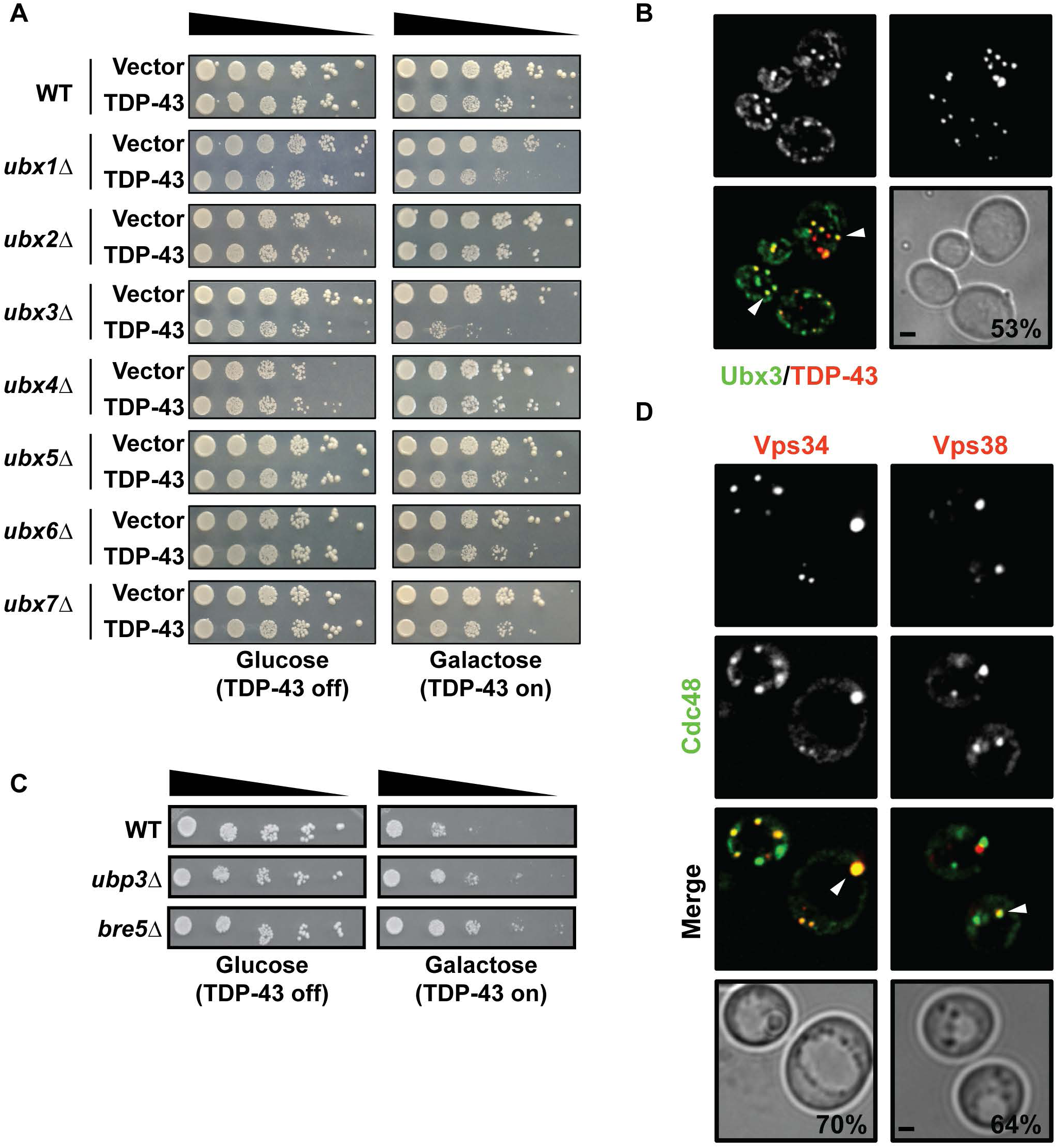
Cdc48 acts in part through endocytosis to regulate TDP-43. (A) Growth assay of WT and indicated isogenic null strains transformed with an empty vector or *GAL1*-regulated TDP-43-GFP plasmid. (B) Ubx3-GFP strain was transformed with a TDP-43-mRuby2 plasmid and images were taken at mid-log phase. Arrowhead indicates Ubx3 and TDP-43 co-localization. Scale bar: 2 µm. (C) Serial dilution growth assay of indicated strains transformed with *GAL1*-regulated TDP-YFP plasmid (D) Cdc48-GFP strain was transformed with an untagged TDP-43 plasmid and either a Vps34-or Vps38-mRuby2 plasmid; Arrowhead indicates Cdc48 co-localization with both Vps34 and Vps38. Scale bar: 2 µm.

We also examined additional gene deletion strains whose genes are either known Cdc48 co-factors, or have been linked to Cdc48 function [49] (Fig 2C). Notably, deletion of *UBP3* and *BRE5* suppressed TDP-43 toxicity. Ubp3 is a deubiquitinase whose activity depends on co-factor Bre5, which also harbors an RNA binding domain. Deubiqutinases play many roles in Ubiquitin metabolism and are generally thought to stabilize proteins by antagonizing Ubiquitin-mediated turnover [50].

Finally, we examined if Cdc48 exhibited any co-localization with endocytic proteins. We focused on subunits in the yeast phosphoinositide 3-kinase (PI3K) complex, namely Vps34 (PI3K itself) and Vps38 (endocytosis-specific PI3K subunit), which aid endocytosis by generating phosphatidylinositol 3-phosphate (PI(3)P) for formation and maturation of early endosomes [51]. We previously showed that both proteins co-localize strongly with cytoplasmic TDP-43-YFP foci in yeast cells [26]. In cells expressing untagged TDP-43, Cdc48 also co-localized well in distinct cytoplasmic foci with Vps34 and Vps38, [51] (Fig. 2D). Together, these findings and the Ubx3 data suggest that Cdc48 regulation of TDP-43 involves endocytic function.

### FUS toxicity and turnover depend on endocytic function and Cdc48

FUS shares many properties in common with TDP-43, including a propensity to form aggregates in ALS and other neurodegenerative diseases including FTLD [2]. A physical interaction between FUS and VCP has also been described [52]. A genetic link between knockdown of Cabeza (FUS ortholog) in *Drosophila*, and altered ter94 expression (VCP ortholog) has also been reported [53]. However, evidence that toxicity of FUS (when expressed at physiological or high levels) or its turnover is regulated by VCP, or indeed by endocytosis in general has not been described. To determine if our findings with TDP-43 [26] were more generalizable to other aggregation prone disease-associated proteins, particularly those linked to ALS, we thus determined if FUS toxicity and turnover could be regulated by endocytosis and Cdc48 as well.

Several findings suggest that FUS, like TDP-43, is regulated by endocytic function in yeast. First, deletion of *VPS15, VPS34* or *VPS38*, which impairs endocytosis, enhanced FUS toxicity (Fig. 3A), similar to our prior work with TDP-43 [26]. Additionally, deletion of several other genes that facilitate maturation of endocytic compartments, including *VPS8* (CORVET complex), *VPS18* (CORVET/HOPS complex) and *VPS21* (Rab5 protein) [54] also exacerbated FUS toxicity in yeast (Fig S1A). In contrast, deletion of core (*ATG1, ATG8*) and selective (*ATG11*) autophagy genes had no effect on FUS toxicity (Fig S1B). Second, deletion of *VPS38*, the PI3K endocytosis-specific subunit, induced FUS accumulation, which can be suppressed by Vps38 re-expression via plasmid (Fig. 3B). Third, FUS foci formation also increased in *vps34Δ* and *vps38Δ* null strains, while showing no increase over WT in *atg8Δ* strains (Fig 3C). Fourth, overexpression of *VPS34* and *VPS38* rescued FUS induced cell toxicity (Fig. 3D). This partly mirrors our prior observation that Vps38 over-expression rescues TDP-43 induced cell toxicity, which correlates with increased endocytic activity [26]. Finally, FUS protein stability showed no stabilization in an *atg1Δ* strain background, but significantly stronger stabilization in a *vps9Δ* (Rab5 GEF), and v*ps34Δ* background (Fig 3E). Interestingly, FUS protein levels were particularly elevated in *vps34Δ* cells versus WT (mirroring Fig 3B), with a modest increase in both *vps9Δ* and *atg1*Δ strains (Fig S1C). Taken together, this suggests that in yeast, endocytic function is a key regulator of FUS toxicity and turnover, while autophagy may play a small role in turnover.

**Fig 3.**
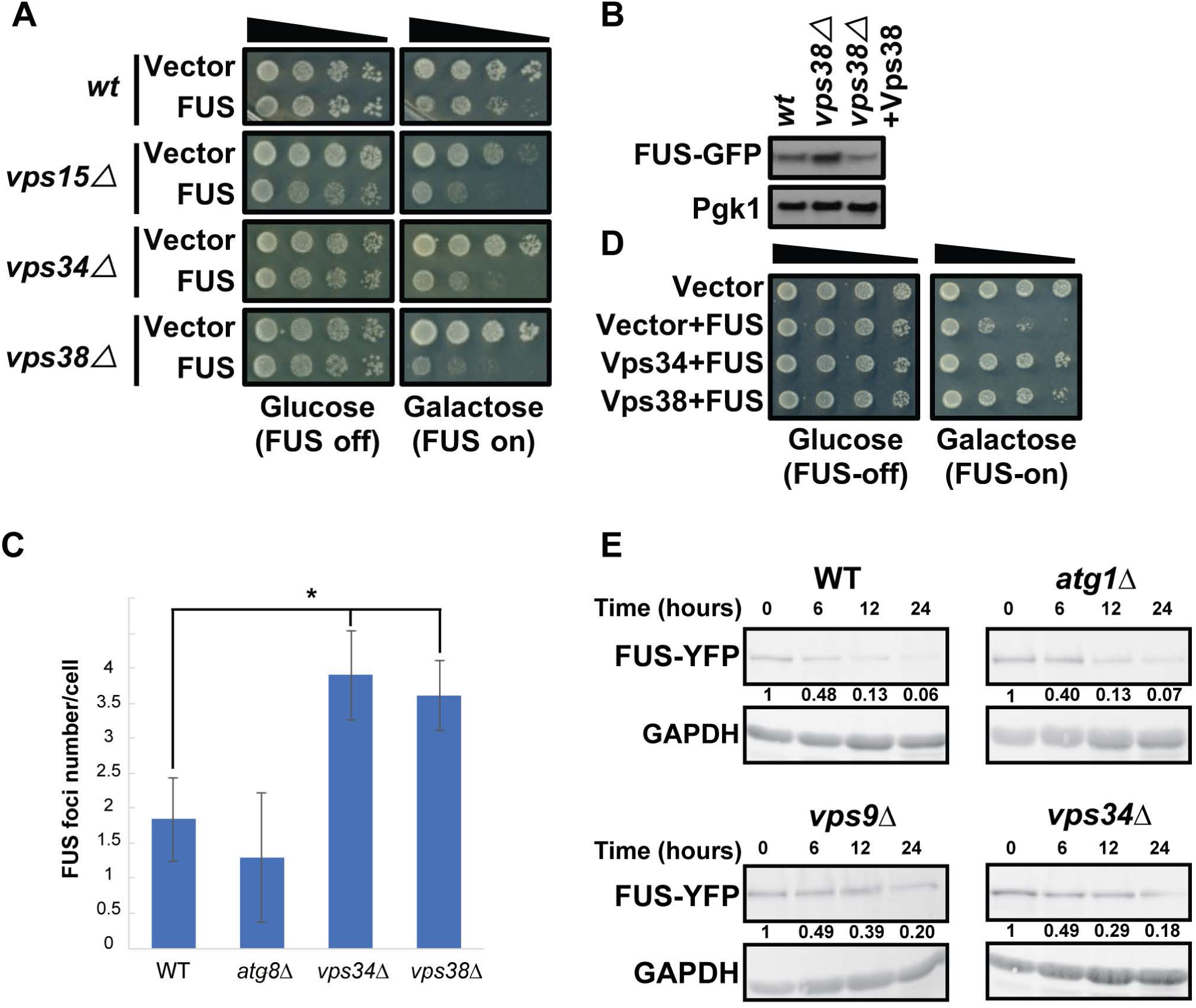
Cdc48 mediates FUS degradation and toxicity via endocytosis. (A) Serial dilution growth assay of indicated strains expressing an empty vector and *GAL1*-regulated FUS-GFP plasmid. (B) Protein level of FUS in the indicated strains; Vps38-mRuby2 plasmid used in rescue. (C) Average number of FUS foci per cell in indicated strains, average of 3 biological replicates. *P < 0.05 by Student’s unpaired two-tailed t test. (D) WT cells with empty vector plasmids, vector plus FUS-YFP plasmid, Vps34-mRuby2 and FUS-YFP plasmids, and Vps38-mRuby2 and FUS-YFP plasmids were tested by growth assay. (E) FUS-YFP protein stability assays following transcriptional shut-off (Glucose addition). FUS levels were quantified following normalization to GAPDH loading control.

In our prior work [26], any endocytosis-related mutant that increased TDP-43 steady state protein levels and/or increased TDP-43 stability, also increased TDP-43 toxicity. Higher transcriptionally-driven expression of TDP-43 also increased toxicity in yeast [26]. However, despite increased TDP-43 protein stability in vacuolar protease mutants [26], TPD-43 toxicity in vacuolar protease mutants does not alter relative to WT cells (Fig S1D). We observe the same phenomenon for FUS (Fig S1E). Thus, FUS and TDP-43 toxicity likely depends on cytoplasmic location and neither proteins are presumably toxic if accumulated within vacuoles.

Finally, we determined whether Cdc48 played a role in regulation of FUS and found similar results to those observed with TDP-43. First, partial and strong inhibition of Cdc48 at 30°C and 35°C respectively enhanced FUS toxicity in the *cdc48-3* strain (Fig S2A). Second, inactivation of Cdc48 at 35°C also lead to a significant increase in FUS protein stability (Fig S2B) Third, of all Ubx null strains examined, only deletion of *UBX3*, which functions in endocytosis [46], clearly increased FUS toxicity (Fig S2C). Interestingly, as with TDP-43, *bre5Δ* and *ubp3Δ* strains also suppressed FUS toxicity (Fig S2D).

In summary, these data suggest that FUS toxicity and turnover, as with TDP-43, is likely regulated in part by a Cdc48-facilitated endocytic process.

### Endocytosis rate is regulated by Cdc48, TDP-43 and FUS

TDP-43 expression in yeast and human cells impairs endocytosis rates, and TDP-43 toxicity and protein accumulation are suppressed by driving endocytosis rates by *VPS38* (yeast) or RAB5 (human cell) over-expression [26]. Having made similar toxicity and protein level observations for FUS in yeast (Fig. 3B and C), we were curious if FUS, like TDP-43, also inhibited endocytosis. We were also curious if defects in endocytosis caused by Cdc48 inactivation using the *cdc48-3* ts allele would be epistatic to, additive or synergistic with effects of TDP-43 or FUS expression.

Yeast endocytosis rates were measured by following uptake of an extracellular dye, FM4-64, which is endocytosed and ultimately stains vacuolar membranes. The faster and more intensely that vacuolar membrane staining occurs, the better endocytosis is functioning [55]. In WT cells, vacuole staining was clearly visible by 5 minutes, and intense by 10 minutes (Fig. 4A, D). Like TDP-43 [26], expression of FUS also impaired endocytosis rates, evident by reduced vacuolar staining over the same time period (Fig 4B-D). Consistent with previous mammalian [31,33] and yeast cell findings [46], inactivation of Cdc48 also impaired endocytosis rates to a similar extent as FUS expression, though not quite as strikingly as TDP-43 expression (Fig. 4A-D). Combining TDP-43 or FUS expression with Cdc48 inactivation lead to a significant additive inhibitory effect on endocytosis (Fig. 4B-D). Taken together, this suggests that both FUS and TDP-43 inhibit endocytosis, in a manner at least partly independent of blocking Cdc48 function.

**Fig 4.**
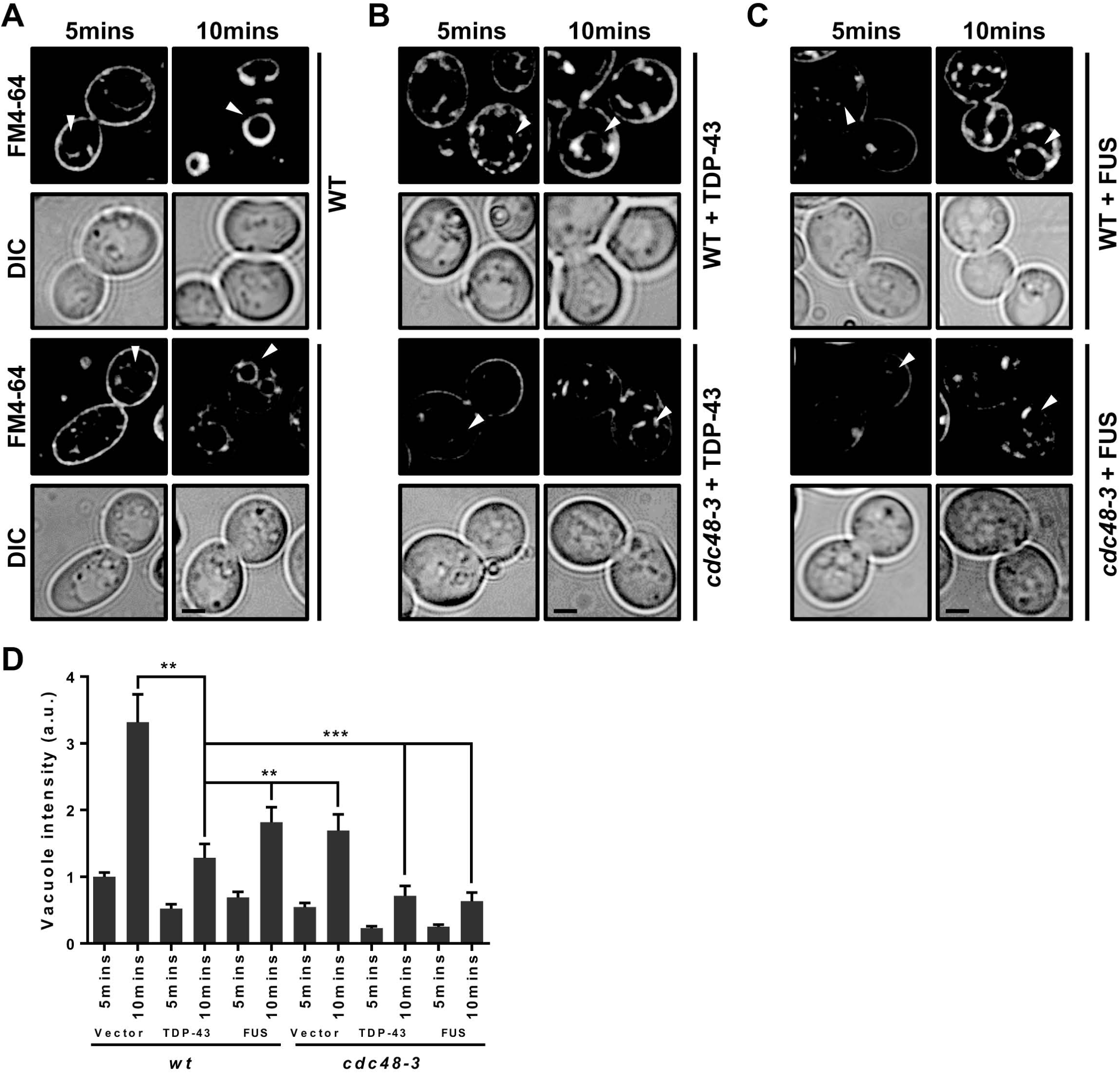
Endocytosis rate is regulated by Cdc48, TDP-43 and FUS. (A) WT and *cdc48-3* cells were cultured at 35°C for 2 hours and then stained with FM4-64 dye (8 uM) for time periods indicated and examined. Arrowhead indicates vacuolar staining. Persistence of plasma membrane staining in *cdc48-3* cells and reduced vacuolar membrane staining relative to WT indicates endocytic defect. (B-C) As in (A) except with additional expression of TDP-43-GFP (B) or FUS-GFP (C). Scale bar = 2 µm. (D) Analysis of vacuolar staining intensity of (A-C). **P < 0.01, ***P < 0.001 by Student’s unpaired two-tailed t test. Data was shown as mean ± s.e.m.

### VCP regulates endocytosis and TDP-43/FUS turnover in mammalian cells

We next turned our focus to VCP in human cells. VCP is a known regulator of endocytosis, localizing to and affecting early endosome size [33], and ultimately affects endocytosis trafficking rates and endocytic substrate turnover in lysosomes [31,33]. First, we examined if VCP localized with Rab5, an early endosomal marker. HEK293A cells transfected with VCP-GFP and Rab5-mRFP constructs exhibited significant co-localization of both proteins (Fig S3A), consistent with prior work in COS7 lines expressing the constitutively active Rab5 Q79L mutant that drives early-endosome enlargement [33]. Next, endocytic uptake of fluorescently-labelled transferrin [56] was performed to measure clathrin-dependent endocytosis rates in WT and VCP knock down cells. Dynasore, a small-molecule inhibitor of dynamin, was used as a positive control of endocytic impairment [57]. As expected, VCP knock down cells exhibited impaired endocytosis, albeit to a lesser degree than in Dynasore-treated cells (Fig S3B); this is consistent with similar data in COS7 cells using Eer1, a VCP inhibitor [33]. These data confirm that VCP positively regulates endocytic function in our cell line model.

We next determined if TDP-43 and FUS steady state levels in HEK293A cells were affected when endocytosis was inhibited. Both dynasore treatment and knock down of VCP increased steady state TDP-43 and FUS levels, mirroring findings in yeast (Fig. 5A). Using TDP-43-GFP lentiviral integrated HEK293A lines [26], we also examined whether VCP inactivation using the specific inhibitor DBeQ altered localization or aggregation of WT TDP-43 or TDP-35 (an N-terminally truncated aggregation prone allele of TDP-43 that mis-localizes to the cytoplasm). Strikingly, in both cases, TDP-43 cytoplasmic aggregation was markedly increased (Figure 5B); whether this reflects loss of VCP functions in endocytosis, or other proteostatic mechanisms, remains unclear. Finally, using immunofluoresence, we examined VCP co-localization with TDP-43 aggregates in frontal cortex tissue from ALS disease and control patients. This revealed that VCP is significantly localized with TDP-43 aggregates in ALS tissue samples, relative to control patients (Fig. 5D, 5E), again mirroring Cdc48 co-localization with TDP-43 in yeast.

**Fig 5.**
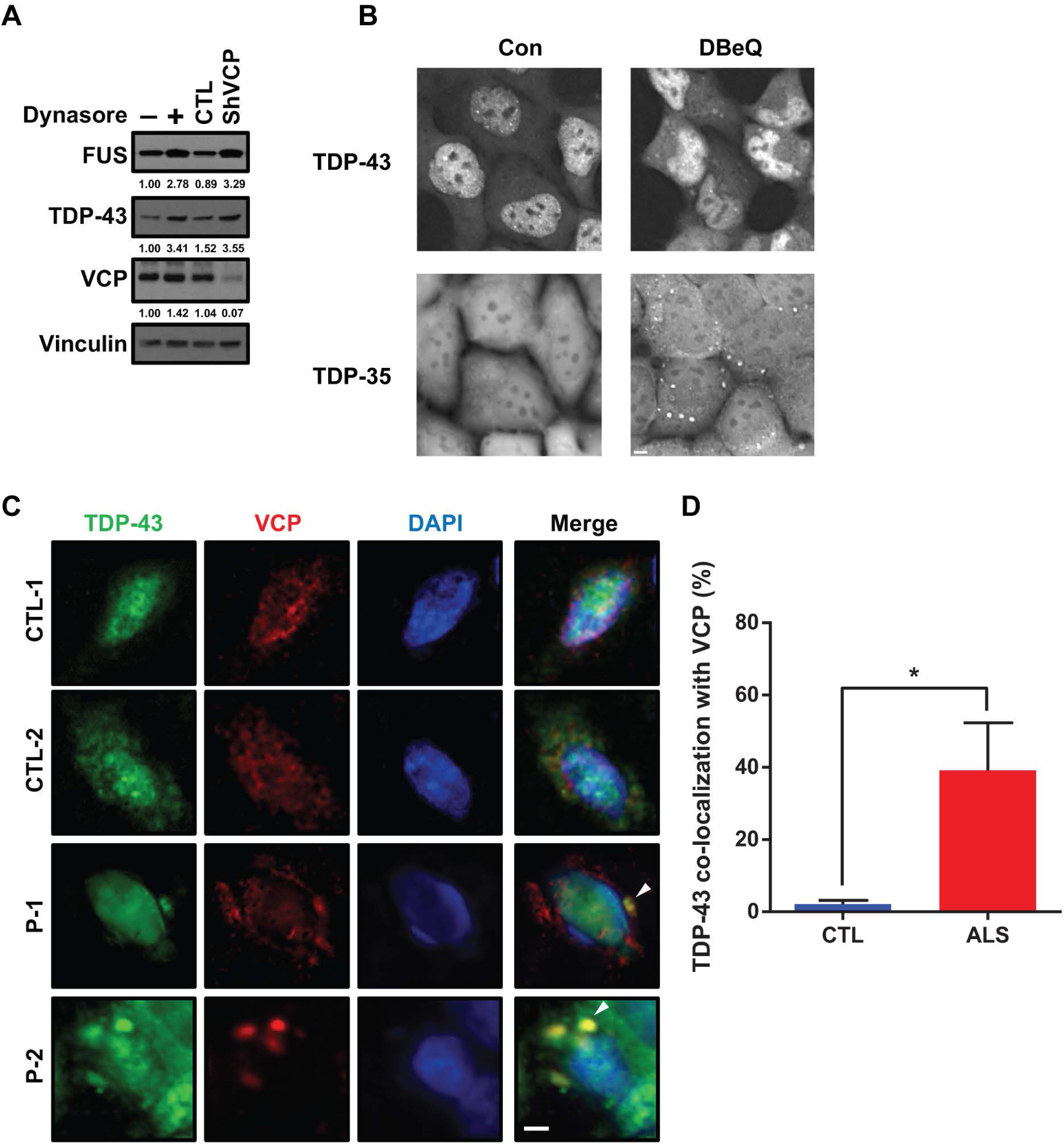
VCP regulates TDP-43 and FUS in mammalian system. (A) HEK 293A cells were either treated with 40µM dynasore for 24 hours or subject to VCP knock down, followed by determination of TDP-43 and FUS protein levels and analyzation. (B) HEK293A cells expressing stably integrated TDP-43-GFP or TDP-35 YFP were examined under normal growth or following DBeQ treatment (10µM, 1hr). Scale bar: 5µm (C-D) Fluorescence immunohistochemistry of control (CTL) and ALS patients (P) frontal cortex tissue with TDP-43, VCP and DAPI and quantification. Arrowhead indicates VCP co-localization with TDP-43. Scale bar: 5 µm. *P < 0.05 by Student’s unpaired to-tailed t test. Data is shown as mean ± s.e.m.

In summary, our data suggests that the previously reported genetic interactions between VCP and TDP-43 [36], and re-localization of TDP-43 to cytoplasmic foci in VCP mutant contexts [32,36,37], may reflect a role for VCP in facilitating cytoplasmic TDP-43 turnover, at least in part via an endocytic mechanism. Our yeast data suggests that Cdc48/VCP may also act similarly on cytoplasmic FUS. These findings are relevant to therapeutic development not only for ALS, but other TDP-43/FUS proteinopathies, including FTLD and particularly IBMPFD, where VCP mutations account for most disease cases.

## Discussion

Several pieces of evidence suggest that Cdc48 and VCP’s effect on TDP-43 and FUS toxicity and turnover may depend on an endocytosis-promoting function. First, inactivation of Cdc48 [46], VCP [33], or expression of either FUS or TDP-43 impair endocytosis (Fig. 4-5). Second, combining expression of either of these aggregation prone proteins with Cdc48-3 inactivation further exacerbates endocytosis inhibition (Fig. 4) and TDP-43/FUS toxicity (Fig. 1F, 3D). Third, TDP-43 and FUS toxicity is increased in *ubx3Δ* strains (Fig. 2A, Fig S2B), which promotes endocytosis [38]. In contrast, deletion of all other Ubx Cdc48 co-factors, including Ubx1 and Ubx2, which function in autophagic and proteasomal turnover pathways [47,48], had no effect on TDP-43 or FUS toxicity. Interestingly, Ubx3 also forms part of a membrane associated E3 ubiquitin ligase complex found throughout the endocytic pathway, but particularly at vacuolar membranes, where it is implicated in ESCRT-dependent turnover of vacuolar membrane transporter proteins in the vacuole itself [51]. Fourth, TDP-43 and FUS shows no increase in cellular toxicity in yeast when autophagy is blocked, but do when endocytosis is inhibited [26] (Fig. 3A, Fig. S1A-B). Fifth, Cdc48 and VCP show strong co-localization with endocytic proteins (Fig. 2D, Fig S3A). Finally, mutations in VCP associated with disease can specifically affect interactions with the endocytosis-promoting cofactor UBXD1 [31], though notably defects in autophagy have also been reported for such mutants [60]. However, this may also reflect defects in endocytosis given the interconnected nature of endocytic and autophagic pathways in human cells [61]. Taken together, these data argue that at least part of the mechanism by which Cdc48 and VCP mediate TDP-43 and FUS protein levels, and their toxicity, is by facilitating turnover by a mechanism dependent on endocytosis activity.

VCP and Cdc48 have been linked to several distinct aspects of endocytic function [31,33]. For example, VCP interacts with and decreases the formation of EEA1 oligomers (early endosome-associated antigen 1). EEA1 promotes fusion of endocytic vesicles into early endosomes, thus increased EEA1 oligomerization may explain an increase in early endosome size that accompanies VCP knockdown or chemical inhibition [33]. VCP inhibition and disease mutations also impair endolysosomal sorting and/or eventual turnover of membrane associated proteins such as epidermal growth factor receptor (EGFR) and Caveolar-1 (CAV-1), which itself functions in caveolar-mediated endocytosis [31][62]. A role for a VCP co-factor, UBXD1, in CAV-1 endolysomal sorting and turnover was also identified [31]. VCP also robustly interacts with clathrin in human cells [63], while in yeast, the Cdc48 cofactor Ubx3 interacts with clathrin and functions in clathrin-mediated endocytosis, as does Cdc48 [46].

We propose that TDP-43 and FUS can be degraded in lysosomes/vacuoles by an endocytosis-dependent trafficking mechanism stimulated by Cdc48/VCP function. A key question is how might Cdc48 and VCP promote TDP-43 and FUS turnover via endocytosis? We propose two general models (Fig 6), which are not mutually exclusive. First, as described above and reviewed elsewhere [28,30], Cdc48/VCP may drive various steps in the endocytic pathway. If TDP-43 and FUS can enter endocytic compartments independently of a physical interaction with Cdc48/VCP, *en route* to clearance in vacuoles/lysosomes, then Cdc48/VCP impairment and resulting endocytic downregulation may indirectly result in TDP-43 and FUS accumulation.

**Fig 6.**
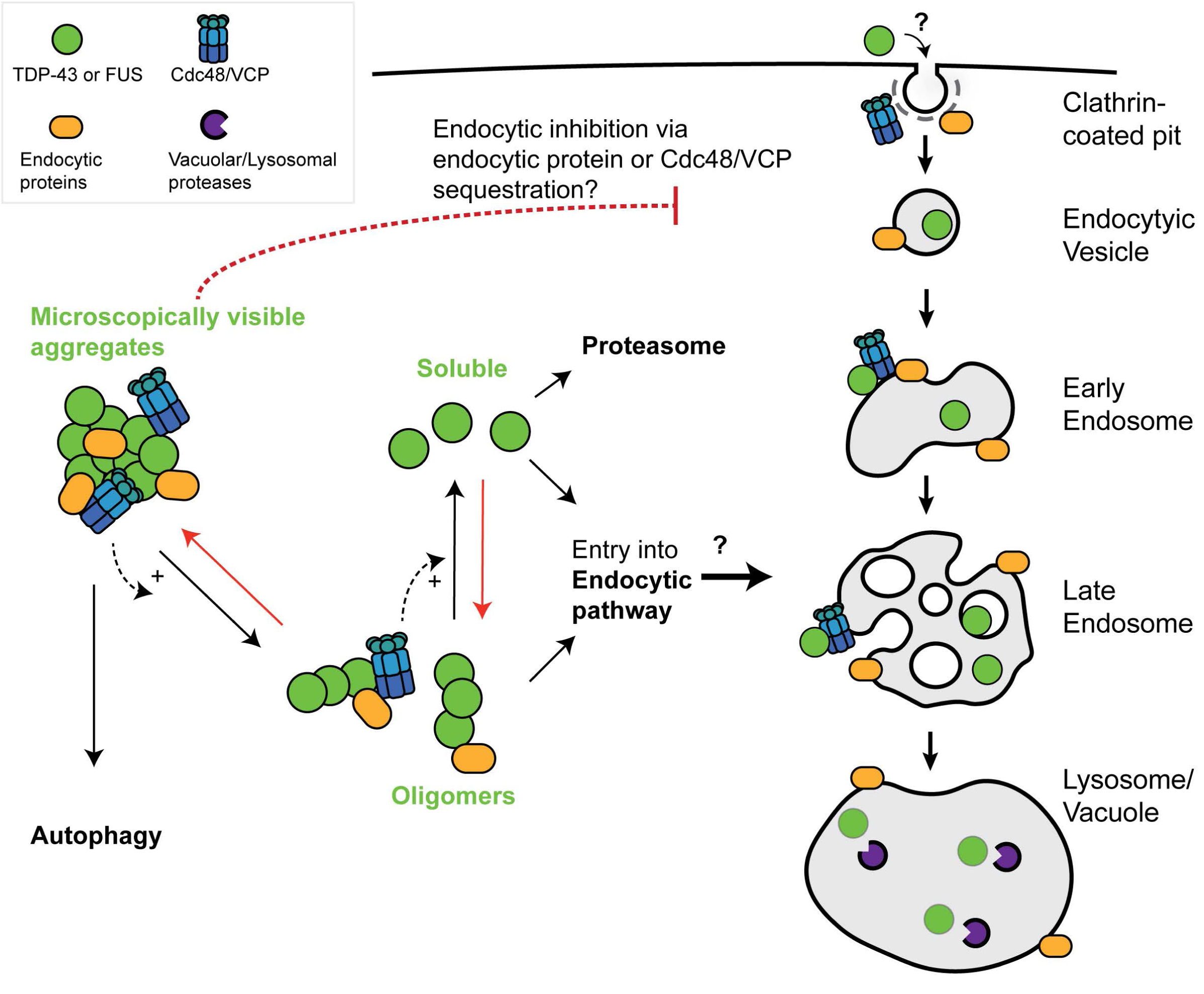
Speculative model of TDP-43 and FUS engagement with endocytosis and Cdc48/VCP function. In yeast, the toxicity, turnover and aggregation of both TDP-43 and FUS depend on endocytosis function (this study and [26]). We suggest that large microscopically visible aggregates of TDP-43 and FUS are most likely cleared from the cytoplasm by autophagy, with oligomeric or soluble forms of TDP-43 and FUS being subject to clearance by endocytic or proteasomal means. If true, given our yeast toxicity data, in which endocytic but not autophagic mutants enhance TDP-43 and FUS toxicity, this would suggest that oligomeric, rather than microscopically visible forms of TDP-43 and FUS are the “toxic” cellular species. Cdc48/VCP, which localizes in microscopically visible TDP-43 or FUS aggregates, may facilitate TDP-43 and FUS conversion into oligomeric and soluble forms. Red arrows broadly represent antagonistic aggregation-promoting processes which may include cellular stress, impaired proteostasis, RNA metabolism defects and perturbation of nuclear-cytoplasmic trafficking, all of which are implicated in ALS pathology. Cdc48/VCP may facilitate entry of TDP-43 and FUS into the endocytic pathway. Finally, sequestration of Cdc48/VCP (and various endocytic proteins) within TDP-43 or FUS aggregates may contribute to the observed inhibition of endocytosis rates caused by TDP-43 or FUS expression, owing to various roles described for Cdc48/VCP in the endocytic pathway. The means by which TDP-43 and FUS enter the endocytic pathway remains mechanistically undefined.

Alternatively, a second possibility is that Cdc48/VCP may directly interact with TDP-43 and FUS aggregates and promote a remodeling event (e.g. aggregate disruption, targeting to endocytic vesicles) that facilitates TDP-43 and FUS trafficking via the endocytic pathway, resulting in their turnover in lysomes/vacuoles. Supporting this, Cdc48 physically interacts with TDP-43 (Fig. 1B), and co-localizes, as does Ubx3, with TDP-43 aggregates (Fig. 1A; Fig. 2B). However, whether these aggregates are active sites of TDP-43 clearance activity or may form part of the mechanism by which TDP-43 impairs endocytosis activity, possibly by sequestration of endocytic factors including Cdc48, remains unclear. Interestingly, TDP-43 and FUS cytoplasmic aggregates are typically ubiquitinated [64,65], thus making them likely substrates for Cdc48/VCP activity. In addition, Cdc48/VCP cofactors may even facilitate TDP-43 or FUS ubiquitination events themselves, given the role of Ubx3 in an E3 ubiquitin ligase complex described previously, and the fact that multiple VCP cofactors interact with E3 ubiquitin ligases [66].

Cdc48/VCP could also promote disassembly and turnover of TDP-43/FUS aggregates by the proteasome or autophagy. However, our prior TDP-43 data in yeast [26], and similar work by the Braun lab [27] argues against a significant role for the proteasome or autophagy in clearing TDP-43 in yeast under normal growth conditions. Similarly, for FUS, no significant impact of impairing autophagy on FUS toxicity was observed (Fig. S1B), though a minor role in turnover was seen (Fig S1C). A key area of future research will be to determine in great mechanistic detail how VCP promotes TDP-43 and FUS turnover in an endocytic-dependent manner.

An intriguing result was the observation that Ubp3 and Bre5 null yeast suppressed TDP-43 and FUS toxicity. The Ubp3/Bre5 complex has been identified as a regulator of numerous vesicular trafficking pathways, including as a negative regulator of mitophagy [67], and a positive effector of ribophagy [68], the cytoplasm to vacuole trafficking pathway [69], and ER-Golgi transport [70]. In yeast, Ubp3 and Bre5 also localize within stress granules [71], and facilitate assembly of stationary phased induced stress granules [72]; indeed Ubp3 and Bre5 are homologs of USP10 and G3BP1, which are also key regulators of mammalian stress granule assembly [73,74]. Thus, a formal possibility (the subject of another work in preparation), is that impaired stress granule assembly may limit TDP-43 aggregation, accumulation and toxicity. Alternatively, loss of Bre5 and Ubp3 function may also reflect altered regulation of various autophagic trafficking pathways or increased turnover of proteins by the proteasome.

Defects in endocytosis have been associated with various neurodegenerative diseases including Alzheimer’s disease and Parkinson’s disease [75–78]. Furthermore, aggregation prone neurodegenerative disease protein clearance via endocytic-dependent means has also been previously suggested. Specifically, α-synuclein degradation in human cell models occurs at least partly by the endolysomal pathway, independent of autophagy and proteasomal turnover [79], while late defects in endocytic trafficking lead to accumulation of polyQ Huntingtin aggregates in yeast Huntingtin models [80]. Several genes associated with ALS onset, including *ALS2, FIG4* and *CHMP2B* also function in endocytosis (discussed in [26]). Additionally, *C9ORF72*, the most commonly mutated gene identified in ALS, also shares DENN domain similarly with Rab GEF proteins, and exhibits co-localization with Rab5. In addition, depletion of *C9ORF72* inhibits endocytosis rates in neuroblastoma cells [81], accelerates degeneration of induced motor-neurons (iMNs) derived from ALS patients, and correlates with endocytic trafficking defects [82]. Strikingly, genetic and pharmacological interventions that enhanced endocytic trafficking in the iMN model, and *C9ORF72* mouse models, also suppressed ALS-associated degeneration phenotypes [82]. Thus, we speculate that endocytic dysfunction may promote accumulation of cytoplasmic TDP-43 and FUS, leading to aggregation. This may be a key part of the pathology of ALS and related degenerative diseases characterized by TDP-43 and/or FUS inclusion bodies, and thus an attractive target for therapeutic development.

## Abbreviations

ALS: Amyotrophic Lateral Sclerosis
CORVET: class C core vacuole/endosome tethering
CHX: Cycloheximide
EGFR: Epidermal growth factor receptor
FTLD: Frontotemporal lobar dementia
FUS: Fused in Sarcoma
HOPS: homotypic fusion and vacuole protein sorting
IBMPFD: Inclusion Body Myopathy with early onset Pagets disease and Frontotemporal Dementia
RRM: RNA Recognition Motif
SG: Stress granule
TDP-43: TAR DNA binding protein 43
VCP: Vasolin containing protein

## Acknowledgements

We thank Southern Arizona VA healthcare agency for providing us with ALS and control patient tissue. This work was supported by start-up funds to JRB from the University of Arizona, NIH RO1-GM1145664, a starter grant (16-IIP-266) from the ALS Association and support from the Frick Foundation.

## Author contributions

JRB conceived the study; JRB and GL designed experiments; GL, AB, FP, AB, EM and AW performed experiments; JRB analyzed data; JRB wrote the manuscript.

## Supplementary Figures

**Fig S1.**
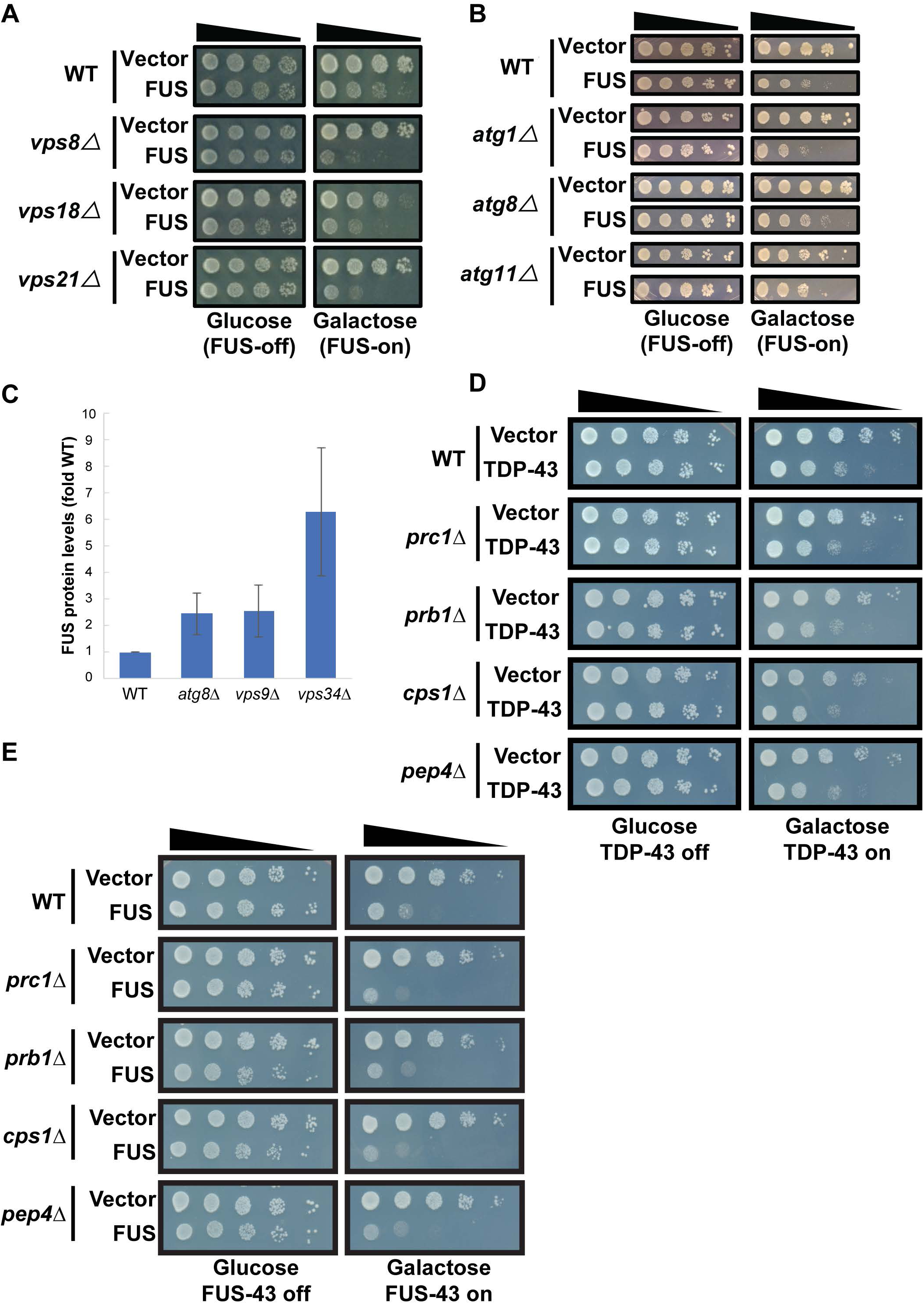
FUS toxicity, aggregation and turnover is regulated by endocytosis in yeast. (A, B) Serial dilution growth assays of endocytosis (A) and autophagy (B) null strains, transformed with a *GAL1*-regulated FUS-YFP plasmid. (C) Quantification of 0hr equal exposure timepoints from FUS stability assays (Figure 3E). Error bars indicate standard deviation from 3 technical replicates. (D-E) Serial dilution assays for indicated strains; performed as in Fig. 2A and Fig. S1A respectively.

**Fig S2:**
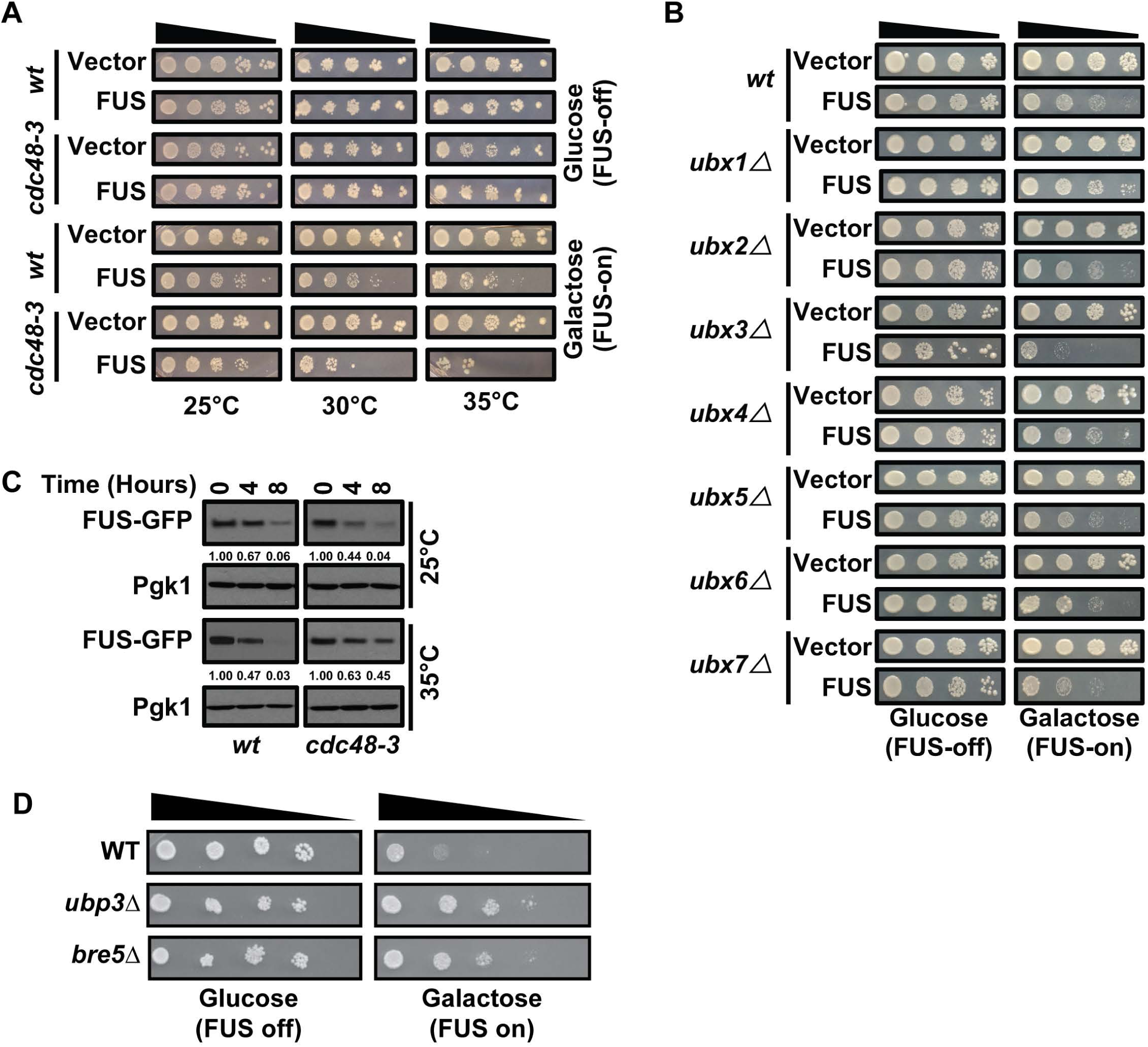
Cdc48 regulates FUS similarly to TDP-43. (A) Serial dilution growth assay of *cdc48-3* strain at permissive (25C), modest (30C) and strongly inhibiting (35C) temperatures expressing an empty vector or *GAL1*-regulated FUS-YFP plasmid. (B) Growth assays in indicated strains expressing empty vector or FUS plasmids. (C) WT and *cdc48-3* strains were treated with 0.2 mg/ml Cycloheximide (CHX) for the indicated time at either 25°C or 35°C. FUS protein stability was quantified following normalization to a PGK1 loading control. (D) Serial dilution growth assay with *GAL1*-regulated FUS plasmid in indicated strains.

**Fig S3:**
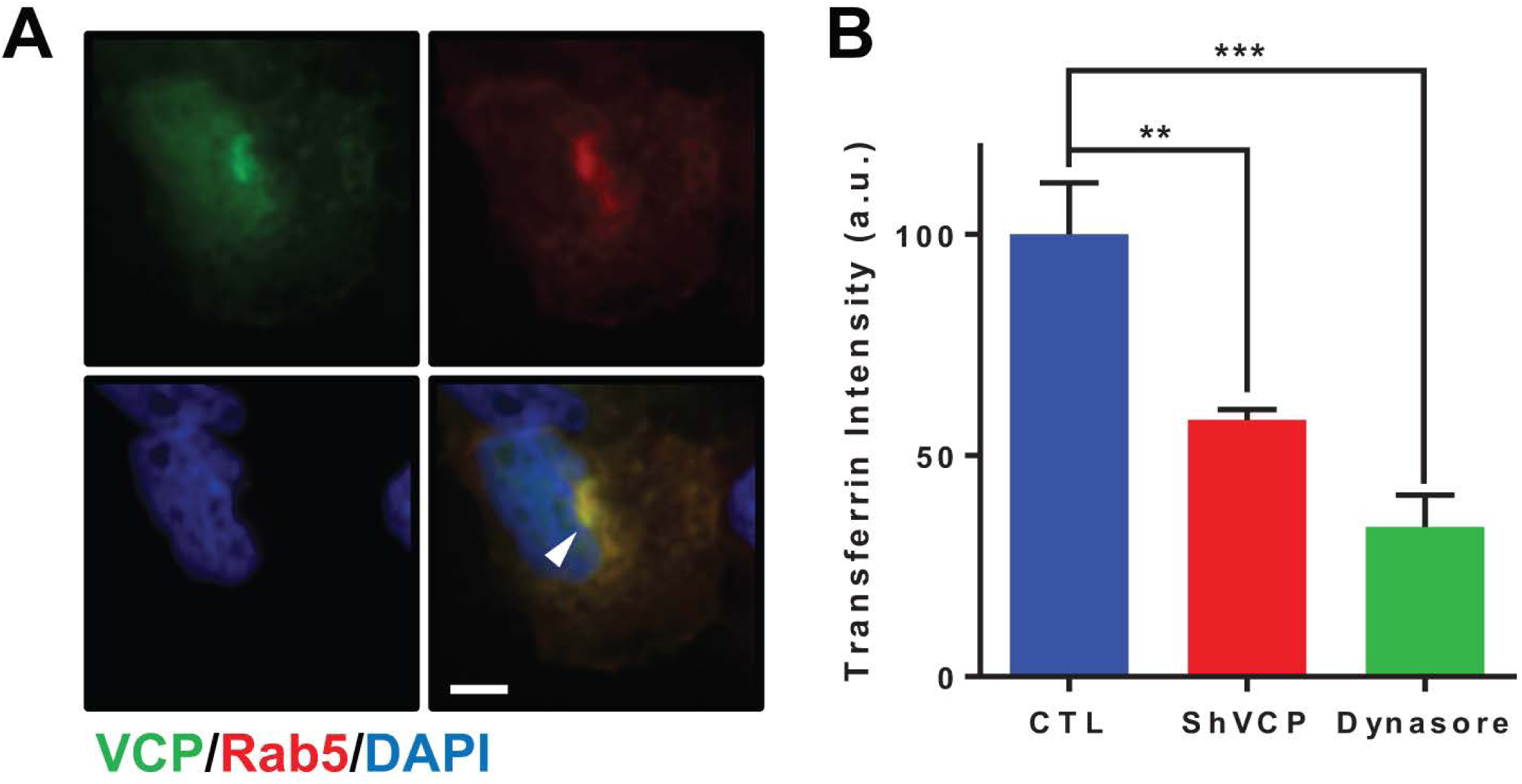
VCP localizes with Rab5 and facilitates endocytosis. (A) HEK 293A cells were transfected with VCP-EGFP and Rab5-mRFP and images were taken. Scale bar = 5 µm. (B) Transferrin uptake assay was performed in control (CTL) and VCP knock down cells. Dynasore (40uM, 24 hours) was utilized as a positive control for endocytosis inhibition. **P < 0.01, ***P < 0.001 by Student’s paired two tailed t test. Data is shown as mean ± s.e.m.

**Table S1.**
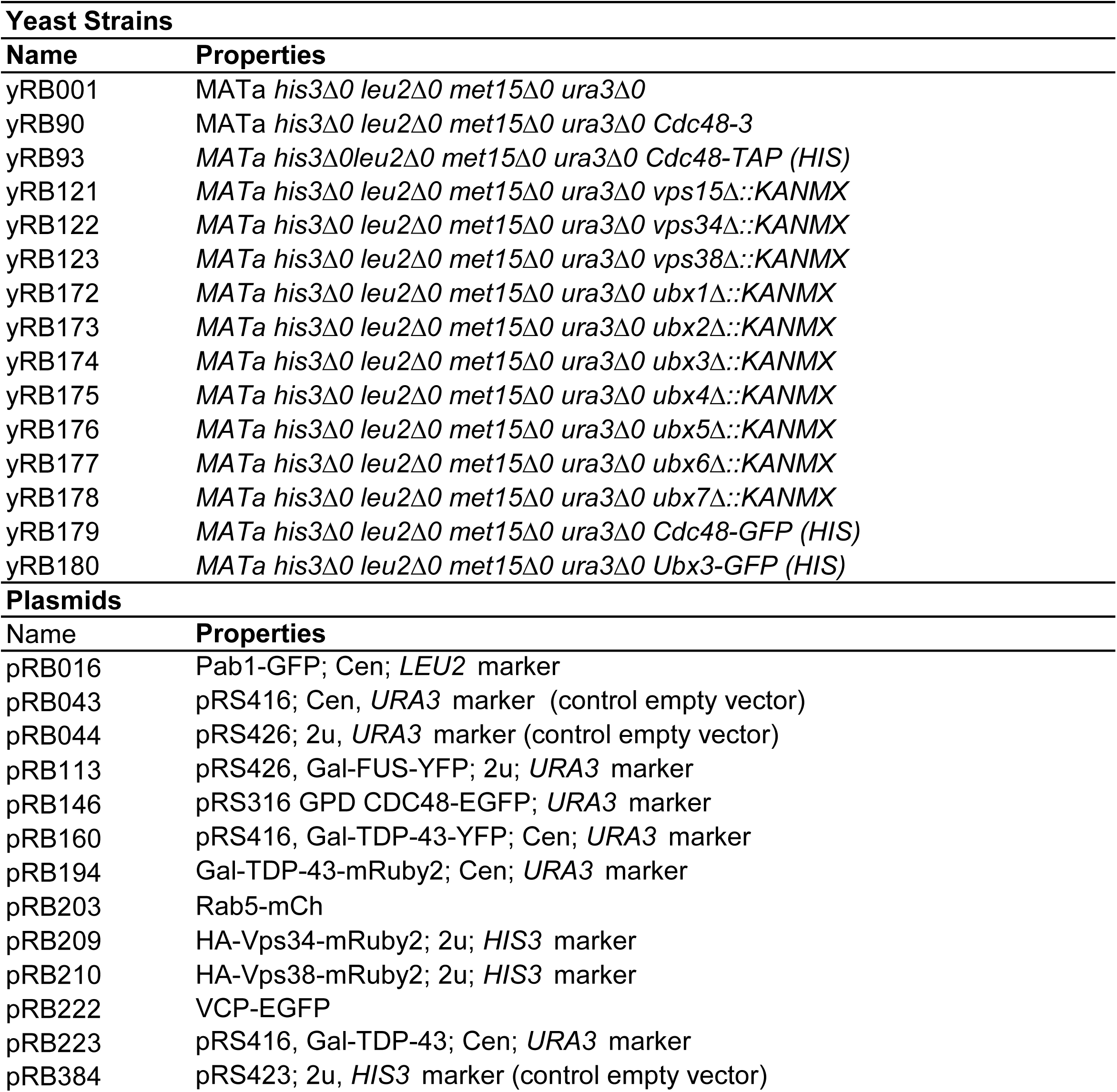
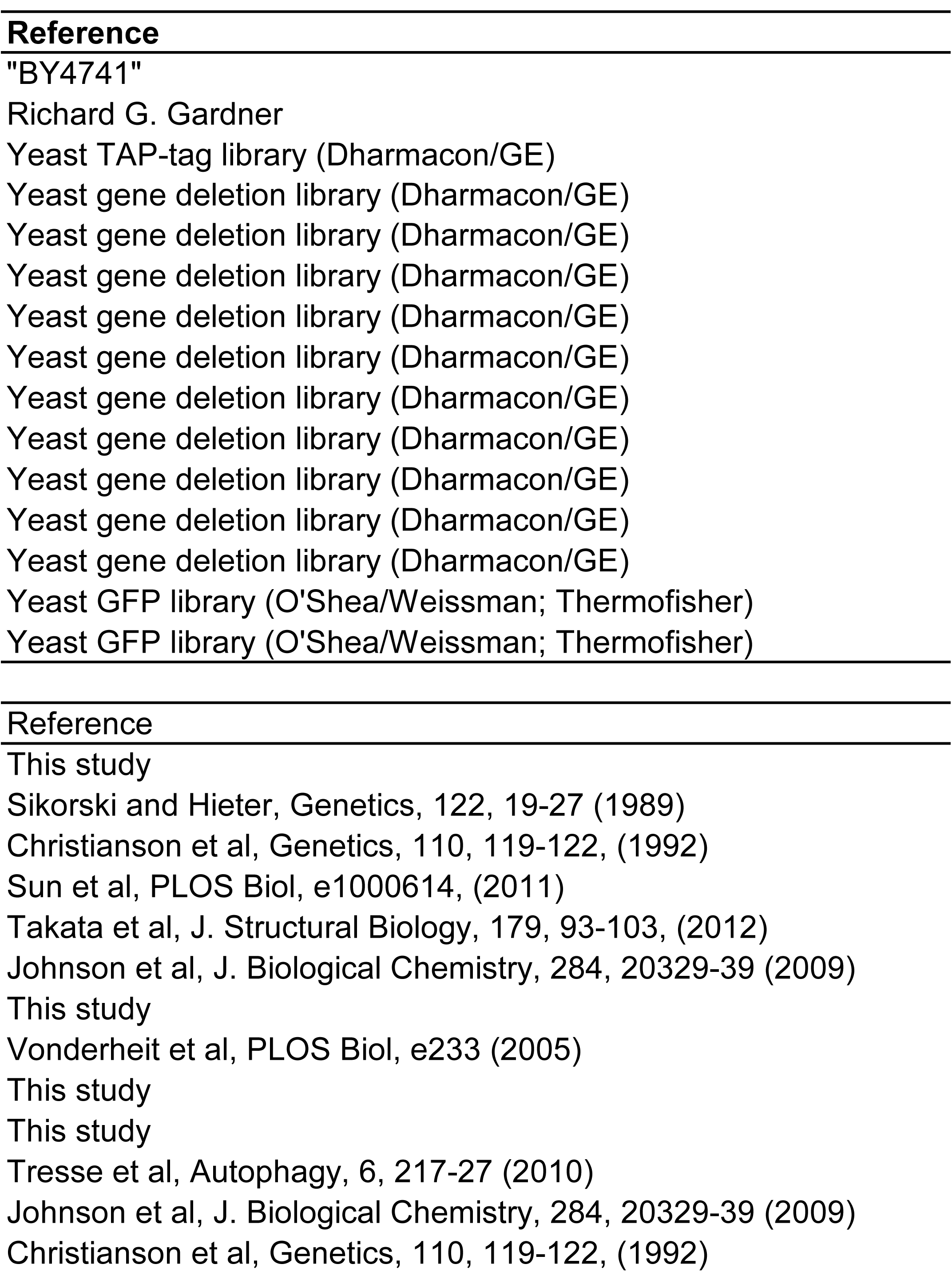
Strains and Plasmids used in this Study.

